# Reading Memory Formation from the Eyes

**DOI:** 10.1101/268490

**Authors:** Anne Bergt, Anne E. Urai, Tobias H. Donner, Lars Schwabe

## Abstract

At any time, we are processing thousands of stimuli, but only few of them will be remembered hours or days later. Is there any way to predict which ones? Here, we show that the pupil response to ongoing stimuli, an indicator of physiological arousal, is a reliable predictor of long-term memory for these stimuli, over at least one day. Pupil dilation was tracked while participants performed visual and auditory encoding tasks. Memory was tested immediately after encoding and 24 hours later. Irrespective of the encoding modality, trial-by-trial variations in pupil dilation predicted which stimuli were recalled in the immediate and 24 hours-delayed tests. These results show that our eyes may provide a window into the formation of long-term memories. Furthermore, our findings underline the important role of central arousal systems in the rapid formation of memories in the brain, possibly by gating synaptic plasticity mechanisms.

---

Our memories define to a large extent who we are. They help us to adapt to current and future realities and without memory any form of education would be unthinkable. Moreover, both the failure to form memories (e.g. in Alzheimer’s disease) and overly strong (emotional) memories (e.g. in posttraumatic stress disorder, PTSD) are clinically highly relevant. Predicting whether, and which, ongoing events will be remembered later on is thus a significant goal of memory research, with considerable implications for various applied contexts.

Recent years have witnessed a revitalized interest in pupil dilation as indicator of phasic changes in central arousal state during cognitive processes ^1–5^. Task-evoked pupil responses reflect phasic activity of neuromodulatory brainstem nuclei controlling arousal. One of these is the locus coeruleus ^6,7^, a brainstem nucleus that provides the major noradrenergic input to the limbic system and neocortex, which is crucial for memory formation ^8^. Thus, the pupil response might provide a window into the making of memories. Yet, so far only very few studies have examined the link of pupil dilation and memory ^9^. In these studies, however, the association of pupil dilation and memory was assessed immediately after encoding and mainly at the group level. Whether the pupil response may forecast the long-term retrieval of individual stimuli remained unknown.

We tested here the hypothesis that the pupil response during encoding of stimuli – and, by inference, phasic elevation of central arousal – predicts trial-by-trial long-term memory formation. In two independent tasks, a visual picture encoding task and an auditory word encoding task, participants encoded either pictures or words while their pupillary responses were tracked with an eye tracker. The use of two tasks allowed us to assess the robustness of the hypothesized predictive value of the pupil response during encoding for subsequent memory. Moreover, because visual stimulation per se leads already to a pupil response, the use of an additional auditory task enabled us to examine the association between pupil dilation during encoding and later memory when any visual artifacts could be ruled out. Memory for the stimuli was tested both immediately after encoding and 24 hours later. Because pupil dilation reflects also emotional arousal ^9,10^ and the well-known memory enhancement for emotional relative to neutral events ^11^ is crucial for several psychopathologies, including PTSD ^12,13^, we included neural and emotionally arousing stimuli to further examine whether pupil dilation may have particular predictive value for emotional memory formation.

Participants’ emotionality ratings during picture encoding confirmed the classification into neutral and negative pictures (mean rating (SEM) for neutral pictures: 0.15 (0.02), for negative pictures: 1.98 (0.05); t(53) = 38.19, p < 0.001, d = 7.35). As expected ^11^, negative pictures were significantly better remembered than neutral pictures, both in the immediate free recall test (t(53) = 12.89, p < 0.001, d = 2.47; Figure 1a) and in the 24 hours-delayed recall (t(53) = 12.39, p < 0.001, d = 2.38; Figure 1b).

**Figure 1:**
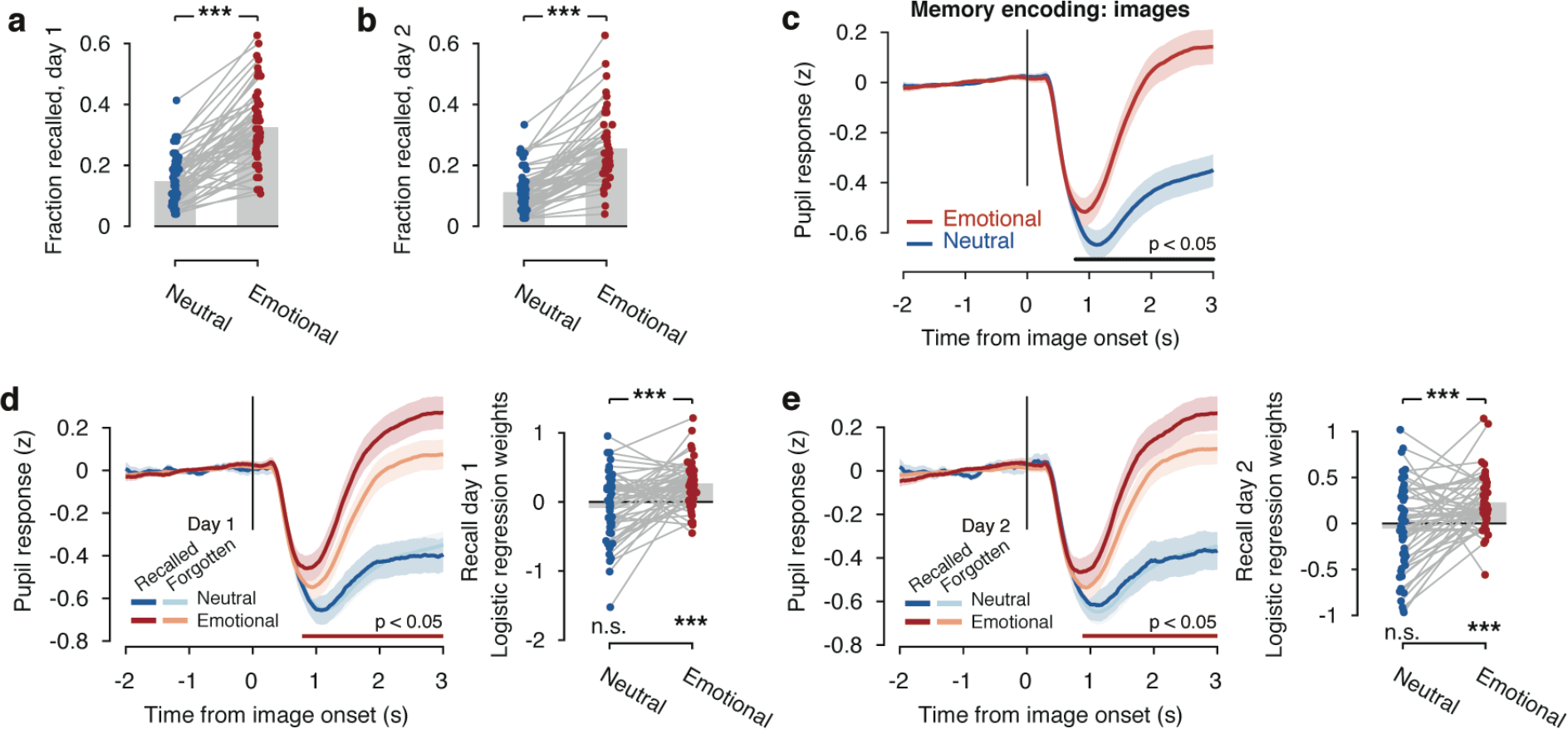
Predictive value of pupil dilation for memory formation in the picture task. (a) Recall performance in the immediate and (b) 24hours-delayed memory, expressed as fraction of recalled pictures, was better for negative than for neutral items. (c) Timecourse of baseline-corrected pupil size in response to picture presentation. After picture onset, the pupil initially constricts in response to the higher contrast images. From 0.78 s after pictures onset, emotion pictures elicit a larger pupil response than neutral pictures. Black line indicates significant timepoints, obtained from a cluster-corrected permutation test. Lines and shaded error regions indicate mean +− s.e.m. (d) Left: Pupil timecourses, separately for emotional and neutral pictures that were recalled or forgotten in the recall test on day 1. Horizontal lines indicate timepoints at which the pupil dilation is significantly different between forgotten and recalled stimuli, as obtained from cluster-corrected permutation test (separately for emotional and neutral stimuli). Right: Individual beta values from logistic regression analyses, indicating that the pupil response during encoding predicts immediate recall of negative but not neutral pictures. (e) Same as panel (d) but for 24hours-delayed recall test. *** two-tailed P < .001.

This emotional memory enhancement was reflected in participants’ pupil responses (Figure 1c). The pupil initially constricted during stimulus presentation, an effect only evident for pictures, not for words (compare with Figure 2c), which is due to the pupil response to the presentation of high-contrast images. This constriction was followed by an evoked dilation, the amplitude of which was modulated by emotional content: Pupils dilated significantly more strongly in response to negative as compared to neutral pictures (t(50) = 11.16, p < 0.001, d = 2.15; Figure 1c).

**Figure 2:**
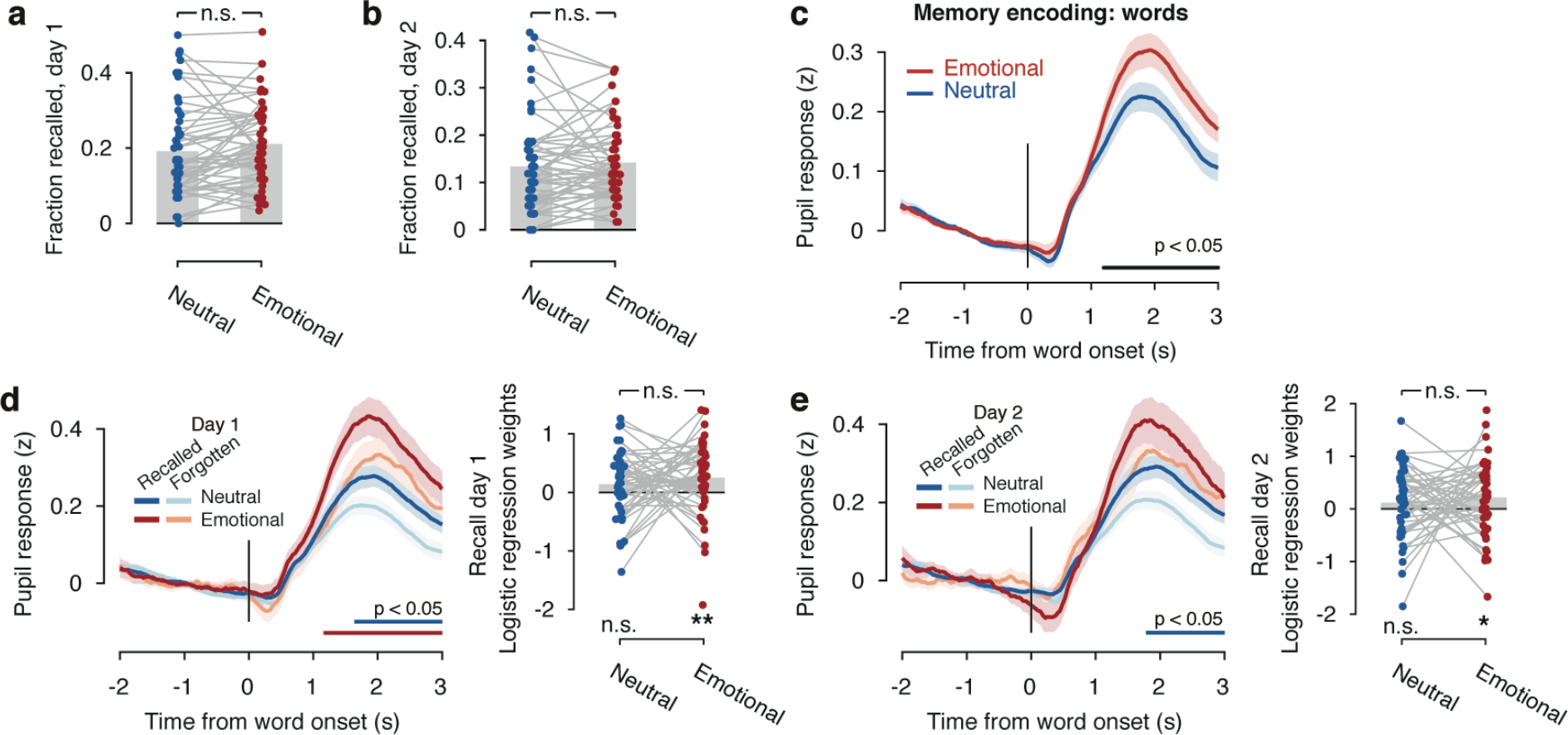
Predictive value of pupil dilation for memory formation in the word task. a) Recall performance in the immediate and (b) 24hours-delayed memory, expressed as fraction of recalled words. (c) Timecourse of baseline-corrected pupil size in response to auditory word presentation. Negative words elicit a significantly larger pupil response than neutral words. Black line indicates significant timepoints, obtained from a cluster-corrected permutation test. Lines and shaded error regions indicate mean +− s.e.m. (d) Left: Pupil timecourses, separately for emotional and neutral words that were recalled or forgotten in the recall test on day 1. Horizontal lines indicate timepoints at which the pupil dilation is significantly different between forgotten and recalled stimuli, as obtained from cluster-corrected permutation test (separately for emotional and neutral stimuli). Right: Individual beta values from logistic regression analyses, indicating that the pupil response during encoding predicts immediate recall of negative but not neutral pictures. (e) Same as panel (d) but for 24hours-delayed recall test. ** two-tailed P < .01, * two-tailed P < .05.

Importantly, pupil dilation during encoding was significantly stronger for items that were subsequently remembered in the immediate free recall test (Figure 1d; main effect subsequent memory: F(1,50) = 9.28, p = 0.004, η_p_^2^ = 0.10) and the 24hours-delayed test (Figure 1e; F(1,50) = 6.62, p = 0.013, η_p_^2^ = 0.09). This effect was driven by the emotionally negative stimuli: Significant interactions of stimulus emotionality and subsequent memory in the immediate (F(1,50) = 10.58, p = 0.002, η_p_^2^ = 0.21) and delayed (F(1,50) = 4.21, p = 0.045, η_p_^2^ = 0.08) recall tests revealed that the pupil response during encoding was larger for remembered than forgotten pictures, when pictures were negative (immediate recall: t(50) = 4.83, p < 0.001, d = 0.97; 24hour-delayed recall: t(50) = 3.81, p < 0.001, d = 0.76) but not when pictures were neutral (immediate recall: t(50) = 0.51, p = 0.610, d = 0.10; 24hour-delayed recall: t(50) = 0.08, p = 0.939, d = 0.02).

These results so far established that pupil responses were larger for negative compared to neutral pictures, and that this pupil response to emotional stimuli was, on average, larger for pictures that were subsequently recalled relative to those that were not. We then set out to determine the predictive power of pupil response for subsequent memory on a trial-by-trial basis, using logistic regression (see Methods). The pupil response during encoding, across items of different emotionality, was a reliable predictor of trial-by-trail memory in the immediate free recall test (average beta-value (SEM): 0.27 (0.04), t-test against 0: t (50) = 6.78, p < .001, d = 1.36) and in the 24hours-delayed test (average beta-value (SEM): 0.26 (0.04), t-test against 0: t (50) = 6.77, p < .001, d = 1.35). When taking the emotionality of the pictures into account, pupil dilation predicted subsequent memory for negative pictures (average beta-value (SEM) for immediate recall: 0.24 (0.05), t-test against 0: t (50) = 5.75, p < .001, d = 1.15; for 24-delayed recall: 0.20 (0.04), t (50) = 5.42, p < .001, d = 1.08) but not for neutral ones (average beta-value (SEM) for immediate recall: −0.09 (0.07), t-test against 0: t (50) = −1.29, p = .211, d = −0.26; for 24-delayed recall: −0.06 (0.06), t (50) = −0.82, p = .418, d = −0.16; Figure 1d and e). These differences in the predictive value of the pupil response for subsequent memory between neutral and negative pictures were statistically significant (immediate recall: t (50) = 4.46, p < .001, d = 0.88; 24hour-delayed recall: t (50) = 3.69, p = .001, d = 0.73).

Thus, the pupil dilation was a reliable predictor of trial-by-trial long-term memory, in particular for emotionally arousing pictures. We replicated these findings in a different stimulus modality, i.e. auditory encoding of words. Recall performance was not reliably different for neutral and negative words, neither immediately after encoding (t(52) = 1.60, p = 0.115, d = 0.31; Figure 2a), nor after 24 hours (t(52) = 0.74, p = 0.462, d = 0.08; Figure 2b). The finding that the emotional modulation of memory was overall lower (if present at all) for the words than for the pictures may be due to fact that the learned words were more abstract than the pictures, in combination with the well-known inferior memory for words relative to pictures ^14^ (see also Figures 1a and b, 2a and b). However, also during the encoding of words the pupil dilation was significantly stronger for negative compared to neutral words (t(48) = 2.72, p = 0.009, d = 0.56; Figure 2c). Note that this emotion-related pupil dilation could not be explained by any differences in visual stimulation as items were presented auditorily.

Again, the pupil dilation was overall stronger for words that were remembered in the immediate (F(1,48) = 23.59, p < 0.001; η_p_^2^ = 0.32) and delayed free recall test (F(1,46) = 10.02, p = 0.003, η_p_^2^ = 0.15) compared to those that were not remembered. This subsequent memory effect was not influenced by the emotionality of the stimuli (subsequent memory × stimulus emotionality for the immediate recall: F(1,48) = 0.09, p = 0.763, η_p_^2^ = 0.00; for the 24hour-delayed recall: F(1,46) = 0.04, p = 0.840, η_p_^2^ = 0.00), suggesting that memory for both neutral and negative words was predicted equally well by the pupil response during encoding, as displayed in Figure 2d and e (left panels).

Again, we further used logistic regression to assess the predictive value of the pupil response during encoding for subsequent memory for words on a trial-by-trial basis. Across neutral and negative items, pupil dilation was a reliable predictor of recall performance in both the immediate (average beta-value (SEM): 0.25 (0.05), t-test against 0: t (48) = 5.00, p < .001, d = 1.01) and in the 24hours-delayed test (average beta-value (SEM): 0.22 (0.06), t-test against 0: t (47) = 4.01, p < .001, d = 0.82). When looking at the neutral and negative words, separately, we obtained that the pupil response predicted the immediate (t-test against 0: t (48) = 2.82, p = .007, d = 0.58) and delayed recall (t-test against 0: t (48) = 2.12, p = .039, d = 0.43) of negative words. For neutral words, the pupil response did not significantly predict immediate or delayed recall (immediate recall: t-test against 0: t (47) = 1.66, p = .103, d = 0.34; 24hours-delayed recall: t-test against 0: t (47) = 1.20, p = .235, d = 0.25). These differences between neutral and negative words, however, were not statistically reliable (immediate recall: t (47) = 1.20, p = .237, d = 0.25; 24hours-delayed recall: t (45) = 0.49, p = .627, d = 0.10; Figure 2h and i).

Our results demonstrate that task-evoked pupil responses predict on a trial-by-trial basis which information will be remembered in the long-run, for at least 24 hours. This effect generalized across visual and auditory encoding tasks, thus allowing us to establish the memory-predictive value of the pupil response across sensory modalities.

Some previous studies have examined the link between pupil dilation and memory ^9,15–18^. Our present findings extend these previous studies in several important ways. First, previous studies have shown that pupil responses, averaged across many trials, differ between memorized and forgotten items. By contrast, we tested here the predictive value of the pupil response at the single-trial-level. Doing so is critical for the prediction of specific memories, as well as for evaluating the utility of pupil dilation as an easily measurable physiological marker of memory formation.

Second, the current study is to the best of our knowledge the first to show that the pupil response predicts whether stimuli will be retained in the long run (for, at least, 24 hours). The delays between encoding phase and memory test were confined to less than 30 minutes in previous work, when memory consolidation, known to take hours ^19^, had (at best) just started. Assessing the stability of pupil-linked memory effects over several delays is important to determine whether the pupil predicts, beyond encoding, also consolidation processes and actual long-term memory and thus to evaluate their real-life behavioral significance. Doing so for two delays a day apart in the current study revealed that pupil dilation predicted the immediate and delayed recall equally well (comparison of beta-weights for immediate and delayed recall in the picture and word encoding tasks: both F < 2.6, both p > .113, both η^2^ < 0.06). This, in turn, showed that pupil responses are reliable predictors of long-term memories and indicates that pupil-linked arousal mechanisms appear to specifically facilitate the encoding of new memories rather than the memory consolidation processes. This is in line with the idea that the phasic release of modulatory neurotransmitters reflected in pupil dilations ^7,20^ help memorize information by gating synaptic plasticity mechanisms in the cerebral cortex ^21^,^22^.

Third, while previous studies used only recognition tests to assess memory, we here assessed both free recall and recognition performance. Free recall provides more insight into the search in, or retrieval from, memory than recognition, while the latter requires merely the comparison of present information to representations in memory. In fact, in our data, pupil dilation significantly predicted only free recall performance (see main figures), but not recognition performance (see supplemental material). This pattern of results might suggest that the arousal reflected in the pupil response aids particularly the search process in memory and less the comparison of information to the internal representation, in line with previous evidence suggesting that free recall is more sensitive to arousal effects than recognition ^23^. Alternatively, the absence of an effect on recognition may also be owing to the excellent (near-ceiling) performance in the recognition test.

Beyond spontaneous or cognitive task-evoked variation in pupil size, the pupil dilates in response to emotionally arousing events ^9,10^. Indeed, we found a modulation of the task-evoked pupil dilations by emotional content, both in the picture and word encoding tasks. In the picture encoding task, pupil dilation predicted memory formation only for stimuli with emotional value, whereas in the word encoding task it did for both, neutral and negative items. This difference might be due to the fact that words were presented auditorily which triggered already a pupil response, corroborating findings showing that tones may lead to pupil dilation and hence promote subsequent memory ^16^.

In conclusion, our findings demonstrate that our eyes may indeed provide a window into the making of long-term memories. So far, subsequent memory paradigms have been used in combination with electroencephalography (EEG) or functional magnetic resonance imaging (fMRI) to identify neural predictors of later memory ^24^. Compared to these complex neuroimaging techniques, pupillometry provides an easily accessible and much cheaper index of memory formation, in particular in the face of recently developed mobile eye-tracking devices. Our data suggest that such devices may be used, for instance in therapeutic or educational settings, to achieve a key goal of memory research, to predict which information will be remembered in the future.

## Methods

### Participants

Fifty-four healthy native speakers of German (age: 18-35 years, M = 25.35 years; 27 women, 27 men) without a history of any neurological or mental disorders participated in this study. All of them reported normal or corrected-to-normal visual acuity and were naïve to the purpose of the experiment. The sample size was based on an a-priori sample size calculation using G*Power ^25^, showing that a sample of 54 participants is required to detect a medium-sized effect of f=0.25 with a power of .95, given an α of .05. Due to technical failure, pupil data were missing for 3 participants in the picture encoding task and for 6 participants in the word encoding task, thus leaving a sample of 51 and 48 participants, respectively, in the corresponding analyses. All participants provided written informed consent and were paid a moderate monetary compensation for participation. The study protocol was approved by the ethics committee of the Faculty of Psychology and Human Movement at the University of Hamburg (approval no. 2016_79).

### Apparatus

The experiment was programmed and presented in MATLAB (The MathWorks, Natick, MA) using the Psychophysics Toolbox ^26^, in combination with the eye-tracking software BeGaze 3.0 (SensoMotoric Systems, SMI). Stimuli were presented on a 24-inch Dell monitor with a resolution of 1920 × 1200 pixels and a refresh rate of 60 Hz. Participants sat in a dimly lit, sound-attenuated room with their head in a chin rest at a distance of 60cm from the screen. Pupil size was monitored in both eyes using a RED250mobile (SMI; sampling rate: 250 Hz). The eye tracker was calibrated applying the 9-point calibration and validation procedure before each of the two encoding tasks.

### Experimental tasks, stimuli and procedure

After their arrival at the lab, participants first completed standard questionnaires to assess their depressive mood, chronic stress level, as well as state and trait anxiety, all of which may affect (emotional) memory processes (see supplementary material). Next, the eye tracker was calibrated and participants performed two encoding tasks: a picture encoding task and a word encoding task. Task order was counterbalanced across participants.

#### Picture encoding task

The stimulus set for the picture encoding task consisted of 150 emotionally neutral and 150 negative pictures taken from the International Affective Picture System (IAPS; ^27^ and other open online sources. Pictures were presented in greyscale and modified in MATLAB so that they all had the same average luminance. During encoding, 75 neutral and 75 negative pictures were randomly chosen from the picture pool and presented in randomized order for 3 seconds at the center of the screen, against a grey background that was equiluminant to the pictures. While encoding the pictures, participants were requested to evaluate the emotionality of the shown picture on a 4-point scale from 0 (“neutral”) to 3 (“very negative”). Between pictures, we presented a grey fixation cross for 3 to 6 seconds.

#### Word encoding task

The stimulus set for the word encoding task consisted of 120 emotionally neutral and 120 negative German nouns. Words were taken from standardized German word data sets ^28,29^. We created audio files for these words with the help of the software Audacity®. During encoding, 60 neutral and 60 negative pictures were randomly chosen from the word pool and presented in randomized order via headphones. While listening to the words, participants looked at a fixation cross shown at the center of the screen with the same grey background as in the picture encoding task. The inter-trial interval between the presentations of words varied between 3 and 6 seconds.

#### Immediate and delayed memory testing

Immediately after each of the two encoding tasks as well as 24 hours after the encoding session, participants performed a free recall test, in which they were asked to report verbally as many of the presented pictures and words, respectively, as possible. The experimenter noted the recalled items on a check list. If it was not entirely clear to which picture a participant was referring to in the free recall test for the picture, he/she was asked to provide more details until the recalled pictured could be clearly identified. There was no time limit for the free recall tests. In order to assess the predictive value of the pupil response during encoding for long-term memory, participants completed a second free recall test 24 hours after encoding. The procedure of this delayed memory test was exactly the same as in the immediate free recall test.

After the 24 hours-delayed free recall test, participants completed also recognition tests for the pictures and words. In these tests, participants saw all pictures and words, respectively, that were presented on the first day and an equal number of novel neutral and negative items in randomized order on a computer screen. Participants were requested to indicate for each item whether it had been presented on day 1 (‘old’) or not (‘new’). For items that were identified as ‘old’, participants were further asked to rate on a scale from 1 (‘not certain’) to 4 (‘very certain’) how confident they were that the items was indeed ‘old’. Because free recall reflects participants’ ability to actually retrieve information better than recognition, free recall appears to be more sensitive to arousal effects than recognition ^23^ and recall and recognition appear to rely on distinct encoding mechanisms ^30^, our analyses focused primarily on the free recall tests. Data for the recognition test are presented in the supplement.

### Pupil data preprocessing and analyses

The pupil data were preprocessed as described in Urai et al. (2017). Missing data and blinks, as detected by the SMI software, were linearly interpolated. We estimated the effect of blinks and saccades on the pupil response through deconvolution, and removed these responses from the data using linear regression. The residual pupil time series were z-scored per run, and resampled to 50 Hz. We segmented the continuous pupil data into epochs corresponding to experimental trials, and baseline-corrected the single-trial data by subtracting the average pupil size in the 2 seconds before stimulus onset. We then defined pupil responses as the average in 1 to 3 s after stimulus onset; this window was chosen to take into account the delay of the pupil response ^2^ and encompass the full presentation duration of the pictures. Statistics on the pupil time course were corrected using cluster-based permutation testing.

### Behavioral data analysis

Data from the picture and word recall tasks on day 1 and 2 were quantified as the fraction of recalled stimuli relative to the number of stimuli presented during encoding. Recall performance for neutral and negative stimuli was subjected to paired t-tests. In order to assess the predictive value of the pupil size during encoding for subsequent memory, we first subjected the data to a subsequent memory analysis, in which we asked whether the pupil size during encoding differed for subsequently remembered and forgotten items. To this end, we subjected the pupil data to an ANOVA with the factors subsequent memory (remembered vs forgotten) and stimulus emotionality (neutral vs. negative).

To analyze on a trial-by-trial basis whether subsequent memory for each individual item can be predicted by pupil dilation during encoding, we employed a logistic regression approach. More specifically, we performed for all participants individual logic regressions estimating the predictive value of the pupil response to an individual item for the subsequent recall of this item. The logistic regression was performed both separately for neutral and negative items and for all items together. The beta-values from these individual logistic regressions were then subjected to t-tests at the group-level to assess whether the beta-values were reliably different from zero and different for neutral and negative items. All reported p-values are two-tailed.

### Data and code availability

All syntax codes used for the analyses as well as al data and materials of the study will be made available by the lead authors upon request.

## Author contributions

L.S. and T.H.D. designed research; A.B. performed research; A.B., A.E.U. and L. S. analyzed data; A.E.U. and L.S. drafted the manuscript, T.H.D. and A.B. provided critical revisions.

## SUPPLEMENTARY MATERIAL

### Supplementary methods

#### Control variables

After their arrival at the lab, participants first completed German versions of the Beck Depression Inventory (BDI; Hautzinger, Bailer, Worall, & Keller, 1994), the Trier Inventory of Chronic Stress (TICS; Schulz, Schlotz, & Becker, 2004), and the State-Trait Anxiety Inventory (STAI; Laux & Spielberger, 1981) to assess their depressive mood, chronic stress level, as well as state and trait anxiety, all of which may affect (emotional) memory processes.

#### Analysis of recognition data

Data from the recognition experiment on day 2 was quantified as the relative fraction of hits and misses, computed on only the stimuli that were previously presented. Additionally, using all the presented (old and new) stimuli, we quantified recognition performance in terms of signal detection-theoretic d’ (Green and Swets, 1966): 
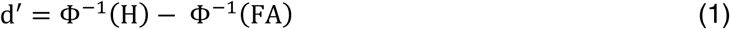
where Φ was the normal cumulative distribution function, H was the fraction of hits and FA the fraction of false alarms. Both H and FA were bounded between 0.001 and 0.999 to allow for computation of d’ in case of near-perfect performance (Stanislaw & Todorov, 1999). This measure gives an unbiased estimate of memory performance. Lastly, we used the average confidence rating for correctly recognized stimuli as a measure of memory strength.

The predictive value of the pupil response during encoding for subsequent recognition memory was analyzed in a comparable manner as for the free recall data (see main text). In brief, pupil data were subjected to an ANOVA to test whether pupil size differed for subsequently correctly detected (‘hit’) and missed neutral and negative items. In addition, logistic regressions testing the predictive value of the pupil response for subsequent recognition performance (hit vs. miss) was performed for all participants and the beta-values form these analyses were analyzed at the group-level.

### Supplementary results

#### Recognition memory performance for pictures

Recognition performance was significantly better for negative than for neutral pictures (Figure S1a and b), as reflected in a higher d’-score (t(53) = 4.97, p < 0.001, d = 0.96) and higher confidence in correctly identified old pictures (t(53) = 6.27, p < 0.001, d = 1.21).

The subsequent memory analysis showed no differences in pupil size during encoding for those pictures that were correctly recognized and those that were not (hits and misses, respectively) 24 hours later (F(1,49) = 0.88, p = 0.352, η_p_^2^ = 0.02; Figure S1c). Whereas there was no difference in pupil dilation for subsequently identified and missed negative pictures (t(1,49) = 0.20, p = 0.839, d = 0.04), there was even a trend for a larger pupil size for subsequently identified vs missed neutral items (t(1,49) = −1.95, p = 0.057, d = −0.39). In line with these findings, the logistic regression analysis showed no evidence for a prediction of subsequent recognition performance by pupil size, neither overall (average beta-value (SEM): 0.02 (0.06), t-test against 0: t (50) = 0.38, p = .704, d = 0.08) nor for negative pictures alone (-ones, as indicated by a higher d’-score (t(52) = 2.42, p = 0.019, d = 0.47) and a higher confidence for correctly identified negative relative to neutral words (t(52) = 2.22, p = 0.031, d 0.07 (0.11), t-test against 0: t (49) = −0.66, p = 0.510, d = −0.13). For neutral pictures, increased pupil size during encoding was even linked to reduced recognition performance 24 hours later (−0.20 (0.09), t-test against 0: t (50) = −2.20, p = 0.032, d = −0.44).

### Recognition memory performance for words

The pattern of results for the word recognition was very similar to the pattern observed for picture recognition. Again, negative words were significantly better recognized than neutral ones, as indicated by a higher d’-score (t(52) = 2.42, p = 0.019, d = 0.47) and a higher confidence for correctly identified negative relative to neutral words (t(52) = 2.22, p = 0.031, d = 0.43; Figure S1d and e). Pupil dilation tended to be higher during encoding of subsequent hits compared to subsequent misses, both overall (F(1,47) = 2.78, p = 0.102, η_p_^2^ = 0.056) and for negative pictures, selectively (t(47) = 1.92, p = 0.06, d = 0.30), whereas there was no such effect for neutral words (t(47) = 0.41, p = 0.683, d = 0.073; (Figure S1f). The logistic regression analysis indicated that the pupil size during encoding was a significant predictor of subsequent word recognition (average beta-value (SEM): 0.15 (0.06), t-test against 0: t (47) = 2.52, p = .015, d = 0.51). When analyzed separately, however, there was a trend for a prediction of negative word recognition by the pupil response (0.14 (0.08), t-test against 0: t (47) = 1.81, p = .077, d = 0.37), whereas there was no such effect for recognition of neutral words (0.06 (0.08), t-test against 0: t (47) = 0.82, p = .414, d = 0.17).

**Supplementary Figure S1.**
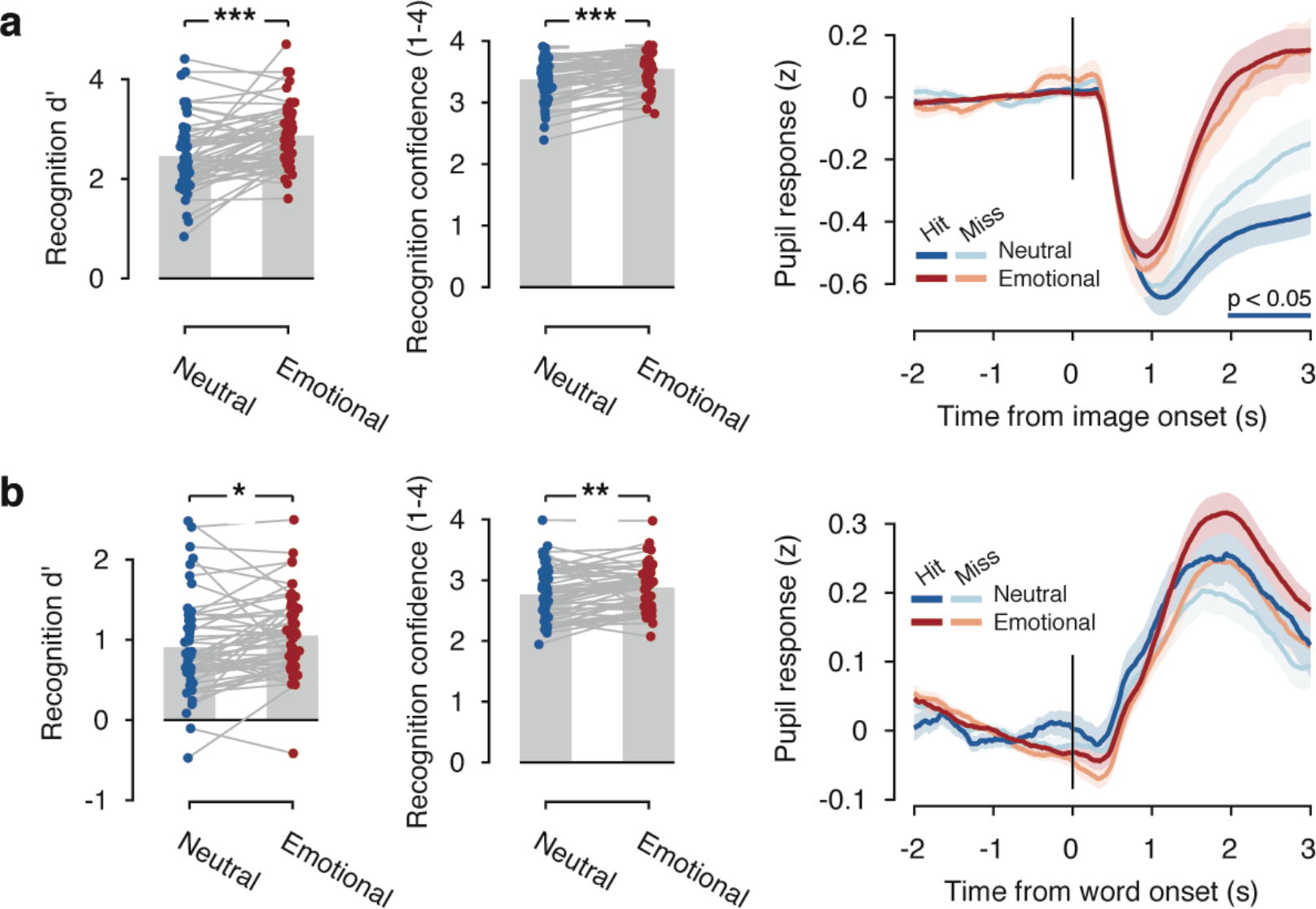
Recognition memory performance in the picture and word tasks. **(a)** Recognition memory expressed as d’ was better for negative than for neutral pictures. (b) Furthermore, participants were more confident in their responses for negative compared to neutral pictures that were correctly identified as ‘old’ (hits). (c) For negative pictures, the pupil dilation during encoding did not differ for later recognized vs. not recognized pictures, whereas for neutral pictures the dilation was even larger for subsequently not recognized pictures. (d-f) Same data but for word encoding task. *** P < .001, ** P < .01, * P < .05.

### Control variables

Participants’ levels of chronic stress, depressive mood, trait and state anxiety were in the normal range of healthy individuals (supplemental Table S1). In order to test whether these parameters were associated with the pupil response during encoding, memory performance and the predictive value of the pupil response for memory, we performed explorative correlational analyses. We obtained negative correlations between both trait anxiety and chronic stress with the predictive value of the pupil dilation for immediate and delayed recall of neutral pictures (all r < -.28; all p < .044; supplemental Table S2). Furthermore, immediate free recall of pictures and both the immediate and delayed free recall of words tended to be negatively correlated with chronic stress level (all r < -.25, all p < .07; supplemental Tables S2 and S3). Although these findings dovetail with earlier reports suggesting impairing effects of chronic stress and trait anxiety on memory (Lupien, McEwen, Gunnar, & Heim, 2009; Pajkossy, Keresztes, & Racsmany, 2017), they need to be interpreted with caution because (i) these correlations would clearly not survive a correction for the number of correlations performed and (ii) it remains unclear why these correlations were observed only for some of the stimuli but not for others.

---

**Supplementary table S1.**
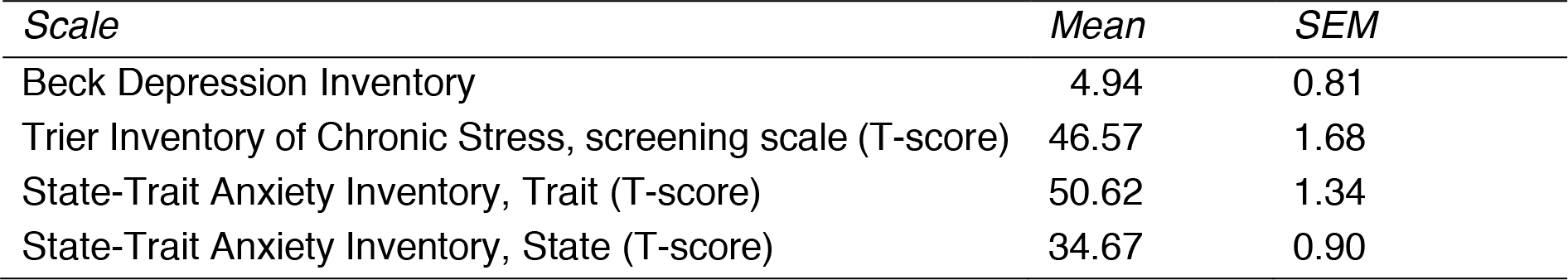
Measures of chronic stress, depressive mood, state and trait anxiety.

**Supplementary table S2.** Correlations of chronic stress, depressive, state and trait anxiety with memory, pupil response and predictive value of pupil response for subsequent recall in the picture encoding task.

|  | Depressive mood | Chronic Stress | Trait anxiety | State anxiety |
| --- | --- | --- | --- | --- |
| Immediate recall, neutral | -0.014<br>*0.920* | -0.268<br>*0.050* | -0.140<br>*0.313* | -0.046<br>*0.741* |
| Immediate recall, negative | -0.094<br>*0.500* | -0.159<br>*0.251* | -0.134<br>*0.336* | -0.022<br>*0.873* |
| Delayed recall, neutral | -0.022<br>*0.875* | -0.164<br>*0.237* | -0.179<br>*0.194* | -0.070<br>*0.614* |
| Delayed recall, negative | -0.032<br>*0.819* | -0.182<br>*0.188* | -0.052<br>*0.709* | 0.033<br>*0.811* |
| Pupil dilation, neutral pictures | 0.025<br>*0.863* | 0.164<br>*0.249* | 0.019<br>*0.894* | 0.010<br>*0.943* |
| Pupil dilation, negative pictures | -0.144<br>*0.313* | -0.005<br>*0.973* | -0.076<br>*0.596* | -0.106<br>*0.460* |
| Prediction immediate recall, negative pictures | -0.80<br>*0.576* | -0.055<br>*0.703* | -0.111<br>*0.439* | 0.022<br>*0.876* |
| Prediction immediate recall, neutral pictures | -0.232<br>*0.102* | -0.283<br>*0.044* | -0.291<br>*0.038* | -0.096<br>*0.501* |
| Prediction delayed recall, negative pictures | 0.044<br>*0.758* | 0.072<br>*0.614* | 0.093<br>*0.515* | 0.170<br>*0.233* |
| Prediction delayed recall, neutral pictures | -0.243<br>*0.086* | -0.363<br>*0.009* | -0.294<br>*0.036* | -0.168<br>*0.238* |

Data show Pearson correlations, two-tailed p-values in italics.

**Supplementary table S3.** Correlations of chronic stress, depressive, state and trait anxiety with memory, pupil response and predictive value of pupil response for subsequent recall in the word encoding task.

|  | Depressive mood | Chronic Stress | Trait anxiety | State anxiety |
| --- | --- | --- | --- | --- |
| Immediate recall, neutral | -0.116<br>*0.409* | -0.262<br>*0.058* | -0.033<br>*0.817* | -0.175<br>*0.210* |
| Immediate recall, negative | -0.180<br>*0.196* | -0.285<br>*0.039* | -0.196<br>*0.161* | -0.261<br>*0.059* |
| Delayed recall, neutral | -0.189<br>*0.175* | -0.439<br>*0.001* | -0.155<br>*0.266* | -0.249<br>*0.072* |
| Delayed recall, negative | -0.074<br>*0.600* | -0.252<br>*0.069* | -0.087<br>*0.534* | -0.132<br>*0.348* |
| Pupil dilation, neutral pictures | -0.128<br>*0.388* | -0.284<br>*0.050* | -0.185<br>*0.207* | -0.093<br>*0.529* |
| Pupil dilation, negative pictures | 0.002<br>*0.992* | -0.075<br>*0.611* | 0.126<br>*0.392* | 0.038<br>*0.798* |
| Prediction immediate recall, negative pictures | -0.014<br>*0.922* | 0.045<br>*0.761* | 0.224<br>*0.126* | 0.202<br>*0.168* |
| Prediction immediate recall, neutral pictures | -0.068<br>*0.645* | 0.115<br>*0.435* | -0.128<br>*0.385* | -0.107<br>*0.470* |
| Prediction delayed recall, negative pictures | 0.040<br>*0.787* | 0.074<br>*0.618* | 0.116<br>*0.430* | 0.239<br>*0.102* |
| Prediction delayed recall, neutral pictures | -0.108<br>*0.473* | 0.115<br>*0.446* | -0.191<br>*0.205* | -0.168<br>*0.264* |

Data show Pearson correlations, two-tailed p-values in italics.

